# Honey bees change the microbiota of pollen

**DOI:** 10.1101/2022.02.21.481367

**Authors:** Alberto Prado, Matthieu Barret, Bernard E. Vaissière, Gloria Torres-Cortes

**Affiliations:** Escuela Nacional de Estudios Superiores, Unidad Juriquilla, UNAM, Querétaro, México; Institut de Recherche en Horticulture et Semences, Agrocampus-Ouest, INRAE, Université d’Angers, SFR4207 QuaSaV, 49071, Beaucouzé, France; Institut national de recherche pour l’agriculture, l’alimentation et l’environnement, UR 406 Abeilles et Environnement, F 84914, Avignon cedex 9, France; Kimitec, Almería, España

**Keywords:** Apis mellifera, corbicular pollen, symbionts

## Abstract

Pollen, as all other plant tissues, harbors different microorganisms. As honey bees (*Apis mellifera*) collect and pack pollen they add regurgitated nectar to moisten and glue the pollen grains, possibly changing the microbial composition. In this study, we compared the microbiota of clean *Brassica napus* pollen with that of bee-worked corbicular pollen. We found that by working pollen, bees increase the bacterial diversity of pollen, by adding honey bee symbionts such as *Bombella, Frischella, Gilliamella* and *Snodgrassella*, bee pathogens as *Spiroplasma* and nectar dwelling *Lactobacillus* to the new pollen microbiota. The bee gut is an important source of inoculum of the corbicular pollen microbiota. We discuss the implications of these findings and propose future research avenues.

## Introduction

Insect visitors shape floral microbiota by acquiring and depositing microorganisms onto flower surfaces during nectar and pollen collection (Keller et al. 2021). The flower microbiota can have important effects on plant (de Vega & Herrera 2012) and pollinator fitness (Adler et al. 2018). Therefore it is important to gain a better understanding of how insects change floral microbial communities. Honey bees are known to introduce bacteria into the nectar while foraging (Aizenberg-Gershtein et al. 2013). In turn, bacteria vectored to the flower change the chemistry of nectar and affect pollinator behavior and plant reproduction (Junker et al. 2014; Rering et al. 2017; Vannette et al. 2013; Vannette & Fukami 2018). However, less is known about the effects of honey bee foraging on the microbiota of pollen. As a reproductive structure, pollen is of great importance for the biological success of plants because its directly linked to reproductive output (i.e. fitness). The few studies that have focused on the microbiota of pollen, have found complex bacterial communities with a high level of species-specificity (Heydenreich et al. 2012; Manirajan et al. 2016). The highly sculptured exine surface of pollen grains present cavities filled with a heterogeneous coating of lipid and proteinaceous components that provide a unique habitat for bacteria (Aleklett et al. 2014; Heslop-Harrison & Heslop-Harrison 1985). Bacteria in the pollen surface can be found alone or in small colonies forming small biofilms (Zasloff 2016). Insect pollinated plants present pollen that has a thick, lipid-rich pollen coat which could be a particular attractive habitat for microbes (Edlund et al. 2004; Aleklett et al. 2014).

The dominant bacterial phyla reported in pollen are *Proteobacteria, Actinobacteria, Acidobacteria* and *Firmicutes* (Heydenreich et al. 2012; Manirajan et al. 2016), at this taxonomic rank no pollen signature compared to other habitats of the plant. However at a finer taxonomic rank, differences in microbiota composition were found between insect- and wind-pollinated tree species, with more similar communities for insect-pollinated species (Manirajan et al. 2016) f. This finding suggests insect vectors affect the microbial composition of pollen by homogenizing the microbial communities across plant species.

Honey bees (*Apis mellifera* L.), one of the most economically important pollinators, harbour a distinct and taxonomically restricted microbiota (Cox-Foster et al. 2007; Martinson et al. 2011) that includes members of the Orbaceae family which are all exclusively honey bee and bumble bee symbionts (Engel et al. 2013; Kwong et al. 2013; Li et al. 2015). The sociality of *Apis* has been proposed as the key driver in symbiont transmission and the maintenance of a consistent gut microbiota (Martinson et al. 2011). Despite the restricted number of species identified in the honey bee gut, symbionts harbour distinct functional capabilities linked to host interaction, biofilm formation, and carbohydrate breakdown (Engel et al. 2012). The Genera previously detected in worker honey bees are *Bartonella, Bifidobacterium, Bombella, Frischella, Gilliamella, Gluconacetobacter, Lactobacillus, Serratia, Simonsiella and Snodgrassella* (Cox-Foster et al. 2007; Engel et al. 2012; Engel et al. 2013; Li et al. 2015; Martinson et al. 2011). The distinctive microbiota of honey bees provides an interesting framework to test the effect of pollinators on the pollen microbiota. In this study we estimated the composition of pollen-associated bacterial communities of *Brassica napus* before and after it has been collected by the domestic honey bee.

## Materials and methods

Detailed methods can be found in Prado et al. (2020). Inside an insect proof tunnel *Brassica napus* L. were sown in the soil. Once in flower, clean pollen was collected by dissecting closed flower buds and separating the anthers. Anthers were left to dehisce for 4 h at room temperature in glass Petri dishes. To recover the pollen, the dried anthers were placed in a steel tea ball and vibrated using a Vibri Vario pollinator. A total of 3 samples of anther collected pollen were recovered. Nectar was collected from the flowers with 2 μl capillary tubes between the base of the anthers and transferred to 2 ml Eppendorf tubes (N=11). Afterwards, a small five-frame honey bee hive was introduced into the tunnel and honey bee foragers were allowed to forage freely on the *B. napus* flowers. Honey bee foragers were captured with a sweeping net and their pollen loads removed and frozen for further analysis (N=3 corbicular pollen). Honey bee foragers were sampled to obtain surface and gut microbial assemblages. Bees were sonicated in 1 ml of PBS buffer with 0.05% Tween 20. After removing the liquid (bee surface samples, N=3), insects were re-suspended in 1 ml of PBS and crushed to recover the microbes inside the bees (gut samples, N=7). All the suspensions were centrifuged (12,000Å∼g, 20 min, 4 °C) and pellets were stored at −20 °C until DNA extraction.

Total DNA extraction of pollen, corbicular pollen, nectar, bee washes and bee homogenates was performed with the PowerSoil DNA isolation kit (MoBio Laboratories) using the manufacturer’s protocol. Amplification, purification and pooling for amplicon library construction were conducted following the protocol described in Barret *et al*. (2015). PCRs were performed with a high-fidelity polymerase (AccuPrime *Taq* DNA polymerase; Invitrogen) using the manufacturer’s protocol and 10 μl of environmental DNA suspension targeting the V4 of the 16S rRNA gene with the PCR primers 515f/806r. After amplicon purification, a second round of amplification was performed with 5 μl of purified amplicons and primers containing the Illumina adaptors and indexes. All amplicons were purified, quantified and pooled in equimolar concentrations. Finally, amplicons libraries were mixed with 10% PhiX control according to Illumina’s protocols. Sequencing runs were performed in this study with MiSeq reagent kit V3 (600 cycles).

### Data Analysis

Primers sequences of fastq files were first cut off using Cutadapt 1.8. Files were then merged and processed with DADA2 v.1.8.0 according to the recommendations of the workflow “DADA2 Pipeline Tutorial”. The workflow was modified in the truncLen parameter to adjust it to the quality of the sequencing run. The 16S rRNA amplicon sequence variants (ASV) resulting from DADA2 were aligned with a naive Bayesian classifier against the Ribosomal Database Project training set 16 database. Data analyses and visualization were done with RStudio v3.3 using the R package *phyloseq* v1.24.2.

## Results

Microbial assemblages of anther collected pollen consisted of *Proteobacteria* (42%), *Actinobacteria* (42%), *Firmicutes* (10%), *Acidobacteria* (4%) and *Bacteroidetes* (2%). These microbial communities were mainly characterised by the Families *Enterobacteriaceae, Pseudomonadaceae* (*Proteobacteria*) and *Microbacteriaceae* (*Actinobacteria*) (Figure 1A). While collecting and packing pollen in their corbicula, honey bees changed the relative proportion of Bacteria phyla and their overall diversity. Corbicular pollen samples contained *Proteobacteria* (40%), *Firmicutes* (21%), *Actinobacteria* (19.7%), *Bacteroidetes* (9.8%), Fusobacteria (2.8%) and *Tenericutes* (1.4%). The change in the microbial assemblages of pollen was partly done by incorporating bee gut-associated symbionts in the Families *Orbaceae* and *Neisseriaceae (Proteobacteria)*, bee pathogens in the Family *Spiroplasmataceae (Tenericutes)*, nectar-associated taxa in the Families *Sphingomondaceae* (*Proteobacteria*) and *Lactobacillaceae* (*Firmicutes*) (Figure 1A). The abundance of Families *Enterobacteriaceae* (*Proteobacteria*) and *Microbacteriaceae (Actinobacteria)* was reduced in corbicular pollen as compared to anther collected pollen. Bee-gut symbionts such as *Bombella, Frischella, Gilliamella* and *Snodgrassella* were completely absent in anther collected pollen but present in corbicular pollen (Figure 1B). ASVs belonging to the Genus *Lactobacillus* present in nectar were also incorporated into pollen samples by bees. Genera such as *Escherichia/Shigella* and *Pseudomonas* that were initially abundant in pollen were reduced in corbicular pollen (Figure 1C).

**Figure 1.**
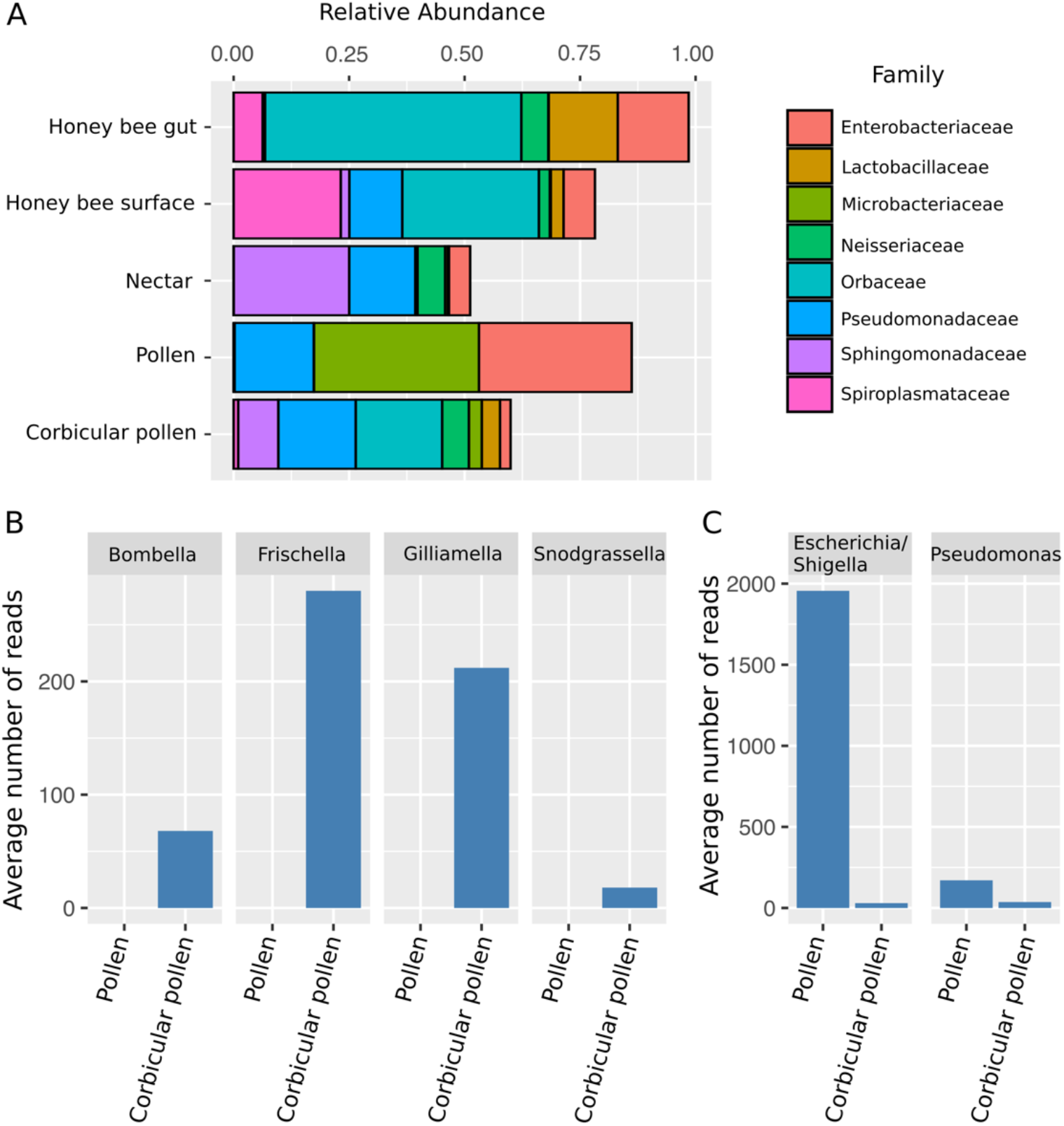
Changes in the microbial composition of pollen. A) Relative abundance of bacterial Families. Only showing Families with more than 3% relative abundance. B) Average number of reads in pollen samples for bee associated taxa. Honeybees introduce *Bombella, Frischella, Gilliamella* and *Snodgrassella* into the pollen. B) Average number of reads in pollen samples for genera that strongly decreased due to honey bee processing.

The microbial assemblages in corbicular pollen were richer than those present in the anther collected pollen with an average of 38 and 15.7 ASVs, respectively (Figure 2A). This enrichment represented 38 additional genera and 60 ASVs, and translated into a higher diversity as expressed by the Shannon index and Faith’s phylogenetic diversity (Figure 2B,C). Of the 71 ASVs present in the corbicular pollen 16.9% are only shared with the bee gut while 7% are only shared with the bee surface, suggesting the potential sources of inoculum. The highest richness was observed in nectar and on the honey bee surface with averages of 54.2 and 53.3 ASVs, respectively. The honey bee gut presented a simpler microbial community with an average of only 19.7 ASVs.

**Figure 2.**
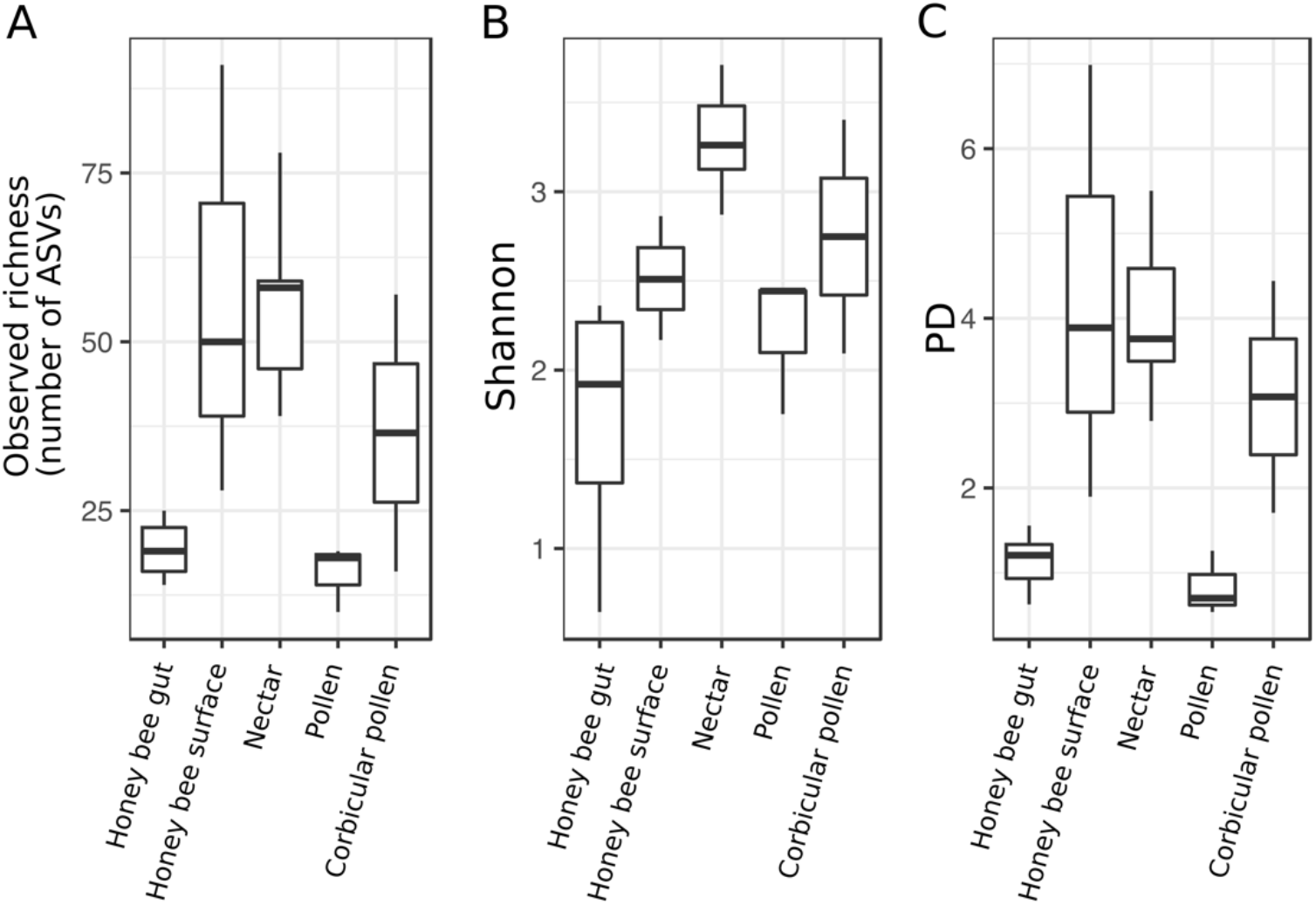
Microbial richness and alpha diversity. Observed richness (a), Shannon diversity index (b) and Faith’s PD phylogenetic diversity (c).

## Discussion

The flower microbiota plays a key role on plant (de Vega & Herrrera 2012) and pollinator fitness (Adler et al. 2018). Therefore its important understand the ecological processes involved in their assembly. This note focuses on the effect of honey bees on the pollen microbiota. Honey bees aggregate pollen grains by adding regurgitated nectar or diluted honey to transport it on the corbiculae of their hind legs. Unsurprisingly, we found that honey bees add bee and nectar-associated taxa to the pollen microbiota. Nonetheless these effects could have important implications for plant reproduction and bee health. The pollen microbiota can have both deleterious and beneficial effects on pollen grain germination on the stigmatic surface. The ubiquitous nectar yeast *Metschnikowia reukauffi* is known to inhibit pollen germination of *Asclepias syriaca* (Eisikowitch et al. 1990), while the nectar dwelling *Acinetobacter* can induce pollen germination of *Eschscholzia californica* (Chistensen et al. 2020). By changing the pollen microbiota honey bees could potentially be modulating pollen germination and hence plant fitness. However, pollen packed in the corbiculae is no longer available for pollination, only pollen grains that remain on other bee surfaces can be deposited on to the stigma. It is unknown if the pollen deposited onto the stigma will suffer similar shifts in its microbial assembly as observed in this study. Nonetheless, honey bees have been shown to change the microbiota of seeds issued from bee pollination by increasing the variability of the microbial communities, and by introducing bee-associated taxa (Prado et al. 2020). Here we show that while collecting pollen bees increase the diversity of pollen bacteria by incorporating nectar and gut-associated taxa. Hence, it is possible that bees’ effect on the microbiota of the seed is a consequence of how honey bees process pollen while foraging. Pollinator behaviour (i.e pollen vs nectar foraging) has been shown to greatly affect the acquisition and deposition of bacteria on to flower surfaces (Russell et al. 2019). If we want to gain a better understanding of the effects of insect pollinators on the plant microbiota, it is crucial to understand the mechanisms involved in the acquisition and deposition of bacteria while insects forage on flowers. Particular attention should be paid to the bacteria vectored to pollen grains and the stigmatic surfaces.

Bee pollen provisions are not consumed immediately after collection, bee larvae usually consume pollen that has been aged. During this process the microbial diversity in pollen provisions of solitary and social bees changes (Graystock et al. 2017; Lozo et al. 2015). Microbes present in aged pollen provisions are central to bee health as a major dietary component (Dharampal et al. 2018) but also as nutritional mutualists allowing bees to digest some of the recalcitrant components of the pollen wall (Engel et al. 2012). Honey bees need to acquire their microbial symbionts twice during their lives, once as larvae and once as adults. Microorganisms acquired by honey bee larvae through food are eliminated during the single defecation prior to pupation, hence worker bees emerging from the cells are usually devoid of microorganisms and re-acquire their microbiota through food and trophallaxis (Gialliam 1997). Therefore the great importance of the presence of bacterial symbionts in the Orbaceae, such as *Frischella* and *Gilliamella*, and the Neisseriaceae, such as *Snodgrassella*, in the pollen provisions (Engel et al. 2012). Our analysis suggests the bee gut is an important source of inoculum of the corbicular pollen microbiota. In conclusion, the way honey bees forage for pollen affects the pollen microbiota with potential consequences on plant reproduction and honey bee health.

## Acknowledgements

We thank Cédric Alaux for his support in planning this experiment, Martial Briand for his help with DADA2 and other bioinformatic analyses. We thank Odile Vilotte, Gilles Aumont and the Agreenskills postdoctoral program for supporting this project. This work was supported by the AgreenSkills+ fellowship programme, which received funding from the EU’s Seventh Framework Programme under grant agreement n° FP7-609398.

